# Motor automaticity in natural keyboard typing

**DOI:** 10.64898/2026.06.04.730281

**Authors:** Rubi Ruopp, Emily A. Williams, Mary Gach, Melissa Baese-Berk, Ian Greenhouse

**Affiliations:** Institute of Neuroscience, University of Oregon, Eugene, OR, USA; Department of Biology, University of Oregon, Eugene, OR, USA; School of Psychology, University of Leeds, Leeds, UK; Department of Human Physiology, University of Oregon, Eugene, OR, USA; Department of Linguistics, University of Chicago, Chicago, IL, USA

**Keywords:** motor control, keyboard typing, automaticity, caching

## Abstract

Certain features of everyday motor skills become automatic while others remain controlled. Here we use a novel keyboard typing task to investigate whether motor automaticity depends on the frequency of naturally learned motor sequences. Participants type five-letter strings that vary in their word and bigram (two-letter sequence) frequency in natural language, allowing us to examine the influence of prior exposure without laboratory training. Novel pseudo word strings are tested as well. We find greater sequence frequency in natural language is associated with faster inter-keypress intervals and lower temporal variability within the sequence. In contrast, latencies to initiate a sequence are slower for novel pseudo-word strings but are otherwise insensitive to natural word frequency. We also find individual differences in inter-keypress speed and variability are robust across frequency levels but are unrelated to conventional measures of typing skill. Our method establishes keyboard typing as a scalable, ethologically valid framework for probing features of a naturally acquired human motor skill. This research will help extend laboratory-based studies of motor sequence learning and sets the stage for future investigations of linguo-motor processes. Moreover, our findings demonstrate which features within naturally acquired motor sequences become automatic and that typing proficiency is not determined solely by automaticity.

**Open practices statement:** All data and analysis code will be made available upon publication at this DOI: https://doi.org/10.5061/dryad.51c59zwrp. None of the reported studies were preregistered.

## Introduction

If you have ever typed an email without actively planning each keystroke, you have experienced motor automaticity. Motor skills, like keyboard typing, are classically described as chunks of motor output strung into flexible, cognitively controlled behavioral sequences (Halford et al., 1998; Lashley, 1951). The individual motor chunks within these skills are considered to be automated: speeded, stereotyped patterns that stabilize with practice and are released from online control (Logan, 1979; Posner & Keele, 1968). Striking the appropriate balance between controlled and automated actions may determine optimal performance of highly practiced behaviors. Expertise is frequently associated with greater automation and may therefore depend on precompiling chunks of motor output for quick and reliable selection and execution.

Nevertheless, it is not clear what features of a highly developed motor skill are automated and what features remain under executive control. Answering these questions can help determine how such skills are mastered and provide insight into the mechanisms that give rise to automaticity.

Automation is a double-edged sword. On the one hand, automated skills are robust to interference from dual tasks that demand increased cognitive effort (Brown & Bennett, 2002; Hassan et al., 2022; Logan, 1979; Poldrack et al., 2005). Thus, automating motor skills frees up cognitive resources. This is beneficial for performing complex tasks, exploring new strategies, and attending to environmental cues (Beilock et al., 2002, 2002; Reingold et al., 2001; Williams & Ford, 2008). On the other hand, automated responses are characterized by inflexible execution of a behavior in a new context, when the response is no longer useful or desired. Because of this feature, habits have long been described in connection, or interchangeably, with automated skills. Habits are defined by behavioral tendencies in which action selection is biased by prior reinforcement (Du et al., 2022; Logan, 1979; Yin & Knowlton, 2006). However, the resulting motor output associated with a habit may not be fast, stereotyped, or low variability – all defining features of automated skills.

Reductions in motor variability are a defining feature of automation. Motor variability has been well researched in the context of novel skill learning (Cardis et al., 2018; Dhawale et al., 2019; Wu et al., 2014; Wulf & Schmidt, 1997) but is seldom studied in the context of a highly practiced motor skill (Marineau et al., 2024). Assessing patterns of variability across levels of frequency of naturally produced sequences can help to identify what features of motor skills are automated and inform possible mechanisms. For example, different neural mechanisms may contribute to motor variability at different levels within a skill, i.e. within chunks or across chunks (Thompson et al., 2019). Linking variability to specific behavioral contexts may help differentiate central from peripheral sources of this variability.

Many behaviors that are self-described as “automatic” take months or years to develop (Nebe et al., 2024). However, most studies of late-stage motor learning and automaticity have used relatively short training periods – typically just days to weeks (Lehéricy et al., 2005; Rosenkranz et al., 2007; Ariani & Diedrichsen, 2019; Yang et al., 2022). While such paradigms are highly effective for studying early-stage skill acquisition and implicit motor learning, this phase of learning is distinct from the stable automaticity that emerges following either explicitly developed expertise or prolonged real-world exposure. This raises an important question: do studies with only a handful of practice sessions truly capture late-stage motor automaticity? Additionally, much of the existing literature relies on well-controlled laboratory paradigms such as the Serial Reaction Time Task (SRTT) and Discrete Sequence Production (DSP) task (Robertson, 2007; Verwey, 2024), which typically involve short sequences and limited training histories and may not reflect the highly practiced, naturalistic motor sequences that characterize everyday motor skills.

A parallel gap exists in research on keyboard typing, as previous studies have not examined typing through the lens of automaticity (Behmer Jr. & Crump, 2016; Logan & Crump, 2009; Pinet et al., 2016; Salthouse, 1984, 1986). Typing on its own warrants further study because typed communication continues to grow as spoken communication shrinks (Pfeifer & Mehl, 2026). The study of keyboard typing can address several of the pitfalls associated with traditional approaches to investigating automated behavior. Typing is ubiquitous, making the behavior well suited to assess naturalistic movement in humans. Traditional keyboard typing was required for 63.3% of US workers in 2019 (Bureau of Labor Statistics, 2019) and digital skills were required for 92% of US jobs in 2023 (Bergson-Shilcock & Taylor, 2023). Typing contains highly stereotyped components at the level of individual keystrokes and combinations of keystrokes (Behmer Jr. & Crump, 2016; Rieger, 2004). Moreover, because it is possible to create completely novel keystroke combinations, keyboard typing allows for the unique opportunity to assess motoric differences between automated and controlled sequences within the same highly practiced, naturally developed skill. While largely unstudied from a neuroscientific perspective, typing has been studied in the fields of psychology (Logan & Crump, 2009; Salthouse, 1984, 1986), linguistics (Pinet et al., 2016; Shulansky & Herrmann, 1977; Will et al., 2006), and cybersecurity (Joyce & Gupta, 1990; Karnan et al., 2011; Stefan et al., 2012). The latter has shown frequently typed strings, such as names and passwords, have highly stable and individualized rhythms of motor output, raising the possibility that typed sequences can be used as reliable identifiers (Joyce & Gupta, 1990).

Here, we designed a novel task to examine motor automaticity that capitalizes on keyboard typing as a naturally developed motor skill. Human participants typed sequences that varied in the amount of prior practice based on their usage in everyday life. We expected more familiar and highly practiced sequences to be more automated than seldom practiced sequences. Using this novel paradigm, we assessed patterns of variability, speed, and reaction time across levels of prior practice and in relation to entirely novel, pseudo-word sequences. We also examined how these metrics differed across scales of action, i.e. bigram (two-letter) and whole word (five-letter) sequences. We found marked differences across sequence frequency levels for both bigrams and words, suggesting natural exposure determines features of motor performance for well-developed, automated skills. The collected metrics were also highly sensitive to individual differences in typing performance. However, these differences did not correspond to a more conventional measure of typing skill, indicating typing skill does not map directly to lower-level features of motor automaticity. Our results are important for understanding the role of automaticity in naturally developed motor skills.

### Materials and Methods Participants and Statistical Power

37 healthy, native English speakers (20 ± 2.9 years of age, 18 female) from the University of Oregon and surrounding community participated in the typing experiment. Fourteen of these subjects self-identified as touch-typists (can type using all fingers without looking at the keys) and seven subjects did not answer the question. Eligibility criteria included no history of neuropsychiatric illness and no use of neuropsychiatric medication. Using G*Power v3.1.9.6 (Faul et al., 2007), this sample size was confirmed to be sufficient to detect differences across bigram and word frequency categories using repeated measures, within-subjects ANOVA testing with a Cohen’s *f* effect size of 0.25, alpha of 0.05, and power of 0.9. The project was approved by the University of Oregon Institutional Review Board, and all participants provided written informed consent.

### Pre-Task Assessments

Prior to the typing task, participants were asked to fill out a short survey regarding their broad motor skill and typing experience (ie. video game play, typed communication in another language) and complete the Edinburgh Handedness Inventory (Oldfield, 1971). Additionally, a subset of 24 participants performed a short typing skill assessment before our task. This assessment, titled Turbo Typing, was used with permission from Dr. Emily A. Williams (Williams, 2024). During this assessment, subjects typed entire sentences, were able to see their typed input, and allowed to backspace. These supplements provided independent assessments of individual typing skill.

### 5-Character Typing Task

Participants completed a typing task built with Psychopy v2022.2.5 (Peirce et al., 2019) during which behavioral keystroke data and high speed (120 FPS) video was collected. Participants were seated in front of a stimulus display monitor and Dell KB212-B QWERTY keyboard. Go- Pro Hero 9 cameras were positioned to collect footage from four planes: aerial, frontal, left stereo, and right stereo (Figure 1a). Video data are not discussed further in this manuscript.

**Figure 1:**
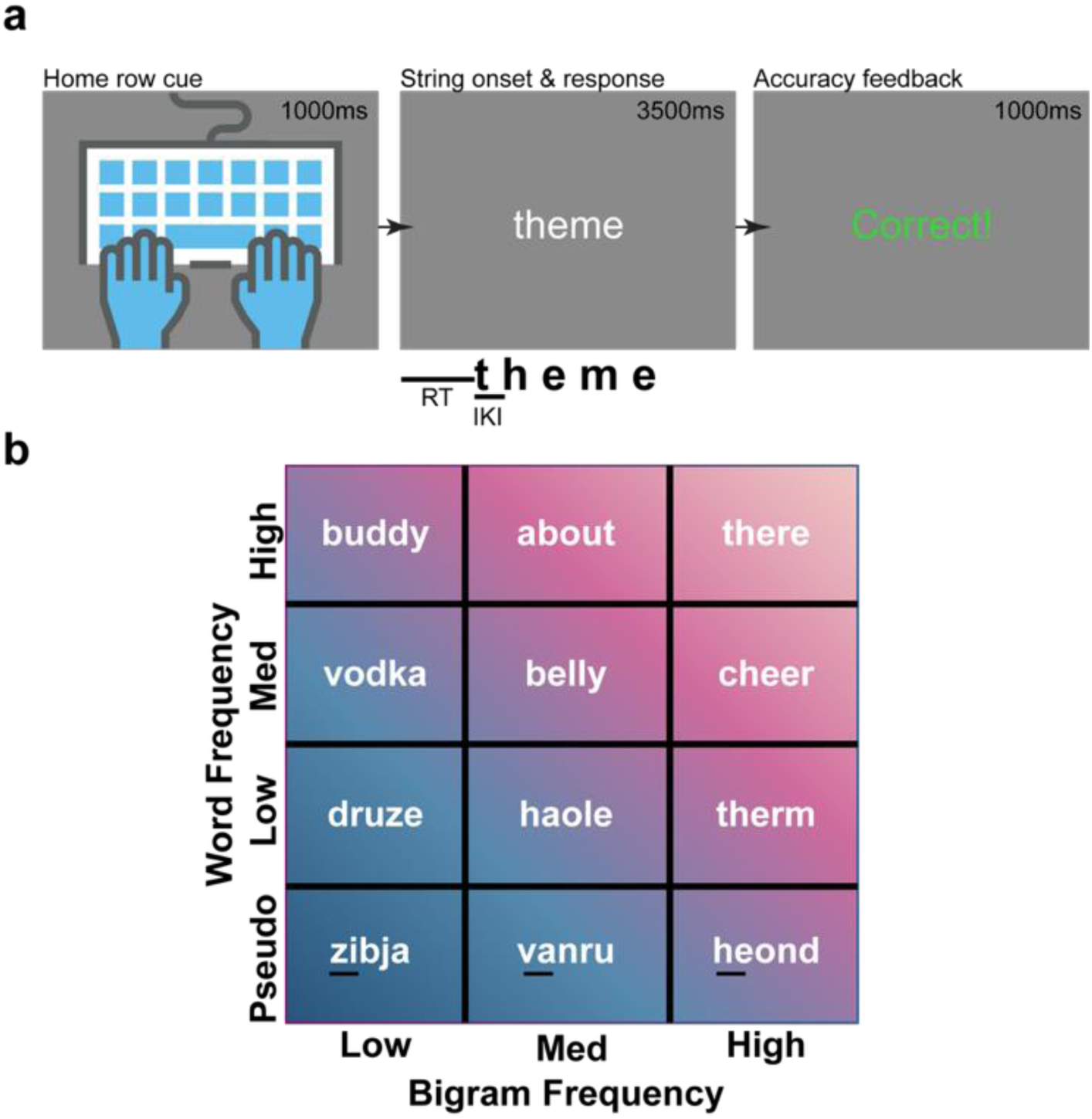
(a) Frames from an example trial of the 5-character typing task. A cue (1000ms) indicated participants should position their hands on the home row of the keyboard. A five - character string was displayed (3500ms), and participants were instructed to type the string as quickly and accurately as possible without backspacing. Accuracy feedback (1000ms) was presented at the end of the trial. (b) Matrix diagram showing example strings from each overlapping word and bigram frequency category. Strings ranged from highly familiar with common subsequences (there) to completely novel with uncommon letter pairings (zibja). Underlined two letter sequences highlight individual bigrams that fall into each of the corresponding frequency categories.

During each trial of the task, participants were seated approximately 70cm in front of an LCD computer monitor (Dell E210H, 60Hz) and cued to type one 5-letter string. Strings were presented in the center of the monitor in white, lowercase, Arial font (108px height), on a gray background, for 3.5s on each task trial. All characters of the string appeared synchronously. A 440Hz tone was played for 100ms through speakers synchronously with stimulus onset to synchronize the video offline. Participants were instructed to type the string as quickly and accurately as possible and, unlike the Turbo Typing assessment, were not provided with online feedback during typing or allowed to backspace. However, delayed feedback immediately following the 3.5s response period and lasting 1s in duration showed the phrase ‘Correct!’ or ‘Oops! That was wrong.’ Trials were separated by a 1s intertrial interval during which an image of hands in the center of a keyboard cued participants to reposition their hands to the home-row. Thus, each trial lasted 5.5s. Participants completed a total of 240 trials administered in two blocks with a short self-paced break (maximum time of 5min) between blocks.

The task was comprised of 24 different strings each repeated 10 times. Strings represented a range of bigram (two letter sequence) frequencies and overall word frequencies in American English (Figure 1b). Eighteen of the strings were words selected from the SUBTLEX corpus (Brysbaert and New, 2009) and categorized based on their SUBTLWF values (frequency of a given word per million words). These words were placed into low (all 0.02), medium (10-19), and high (103-1348) word frequency categories, each containing six words. A category of six word-adjacent nonsense strings was also included to provide entirely unfamiliar words, all representing a SUBLTWF value of zero. Within each of these word frequency categories, strings were also selected to represent a range of average bigram frequencies (mean frequency of all bigrams within each string). Although high frequency words are inherently composed of high frequency bigrams, this categorization allowed us to control, as much as possible, for the influence of bigram frequency and to isolate features of automaticity and variability that are specific to either the sub-sequence or whole-sequence level. Average bigram frequencies were based on total bigram counts (number of occurrences within the entire corpus) calculated from the Project Gutenberg corpus (Behmer Jr. & Crump, 2016) and categorized into low (54063-797899), medium (3221922-3464632), and high (10008813-16848901) groupings, with a total of eight strings per category. Bigrams within strings were also divided into low (36-514720, 26 total bigrams), medium (560130-4235407, 31 total bigrams), and high (4724108-22288309, 16 total bigrams) frequency categories based on total bigram counts.

Strings with higher word and bigram frequencies are assumed to be more practiced, and therefore more automated sequences, while strings with lower word and bigram frequencies are assumed to be less practiced, and therefore more cognitively controlled, sequences. Choosing strings based on their linguistic features allowed real world occurrences to dictate groupings in our task design. We presumed these real-world frequencies correspond to exposure and typing experience within our sample and that more experience typing a particular sequence would result in more automatic motor execution. Additionally, our selection process isolated levels of prior exposure at the string and sub-string levels, allowing for independent analyses for strings and bigrams.

### Metrics of Interest

During the task, all keypress times were recorded for further analysis. The primary metric of interest was interkeypress interval (IKI), the time between the start of two consecutive key presses. IKI was calculated for each bigram of each word across the task. When all IKIs from a string are combined, they effectively evaluate the typing rhythm of a particular word. This collective IKI measure has been shown to be sensitive to individual differences (Joyce & Gupta, 1990). IKIs longer than one second were excluded from further analyses.

For each participant, mean IKI, standard deviation of IKI (SD IKI), and coefficient of variation of IKI (CV IKI) were calculated within (frequency-specific) and across (global) all bigram and word frequency categories. We calculated the mean change in IKI across all consecutive correct trials (mean ΔIKI) separately for each bigram within each word to assess changes in IKI across the course of the task.

Time to initial keypress served as a measure of reaction time (RT) and was calculated for each string within the task. Mean, SD, and CV RT were calculated across word frequency categories at the individual and averaged at the group level. The average change in RT between consecutive trials (mean ΔRT) was measured for each string to determine how time to initial keypress changed over the course of repetitions. These metrics were not calculated across bigram frequency categories, as bigram frequency across first keypress was not controlled.

Error was defined as any mistake made while typing a string or bigram. This definition includes errors of subtraction (missing a letter), addition (adding a letter), and transposition (swapping two letters). Individual total error counts were summed for each bigram and word frequency category. Group level averages were also calculated.

During the pre-task Turbo Typing assessment, baseline typing skill of each participant was determined using the Inverse Efficiency Score (IES) – mean time to each keypress (effectively mean RT+IKI) divided by the percent of correct responses on the character level (one incorrect keypress = 1 error) (Townsend & Ashby, 1983). IES was calculated for each trial and then averaged across trials to generate one mean IES for each participant. Global mean, SD, and CV IKI were also calculated using the Turbo Typing data. As the assessment prompted participants to type full sentences with uppercase letters, spaces, and punctuation, these metrics were calculated from only alphabetic keypresses to be comparable with IKI metrics collected during our 5-character typing task.

### Statistical Analyses

All statistical analyses were performed in Jupyter Lab v4.6.4 with Python v3.12.2 and IPython v8.24.0. One-way repeated measures ANOVA tests performed using the Pingouin 0.5.5 ‘rm_anova’ command compared metrics of interest presented in Table 1 across each of the three bigram frequency categories and/or four word-frequency categories. These tests determined whether metrics differed significantly across levels of automation/frequency.

**Table 1:**
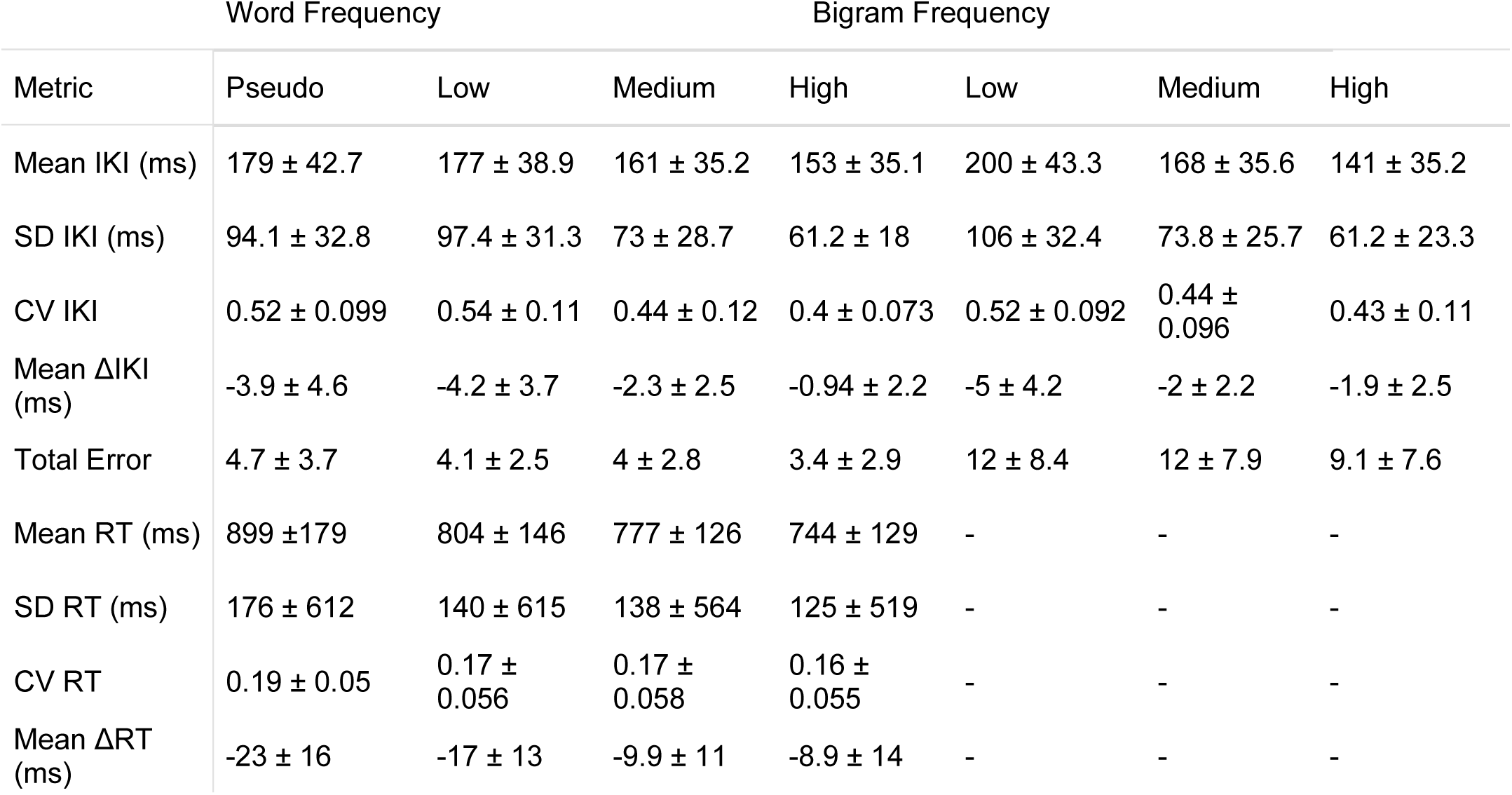
Group level means and standard deviations for metrics collected during the novel typing task. Mean ΔIKI is the average change in IKI between consecutive repetitions of the same bigram embedded within the same string. Mean ΔRT is the average change in RT between consecutive repetitions of the same word. IKI = interkeypress interval.

Pairwise post hoc comparisons were performed using the Pingouin 0.5.5 ‘pairwise_tests’ function and were Bonferroni corrected for multiple comparisons. An alpha value of .05 was used to determine significance. To assess whether individual differences were consistent across categories, Pearson correlation testing performed using the SciPy 1.13.0 ‘pearsonr’ command compared individual mean, SD, and CV IKI values across all bigram and word frequency categories. Additionally, individual mean IES and global mean, SD, and CV IKI scores calculated from the pre-task Turbo Typing assessment were compared with global mean, SD, and CV IKI scores from the 5-character string task. Bonferroni corrections were used for all Pearson correlations.

### Results Speed and variability across levels of prior exposure

To determine how typing speed changed across levels of prior sequence exposure, we assessed Mean interkeypress interval (IKI) across bigram (low, medium, high) and word (pseudo, low, medium, high) frequency categories (Figure 2a, top row). Within participants, Mean IKI decreased with increasing bigram frequency [(F(2,72)=214.9, p=9.2e-26, η^2^_G_=.30]. Post-hoc t-tests showed Mean IKIs for low (200±43ms), medium (168±36ms), and high frequency bigrams (141±35ms) were all significantly different from each other [p’s<.01]. This result indicates the production of naturally acquired two-key motor sequences is sensitive to prior exposure. Mean IKI also showed a significant main effect across word frequency categories [(F(3,108)=37.9, p=4.0e-13, η^2^_G_=.08]. However, post-hoc t-tests were only significant when comparing pseudo (179±43ms) and low frequency words (177±39ms) to high frequency words (153±35ms) [p’s<.05].

**Figure 2:**
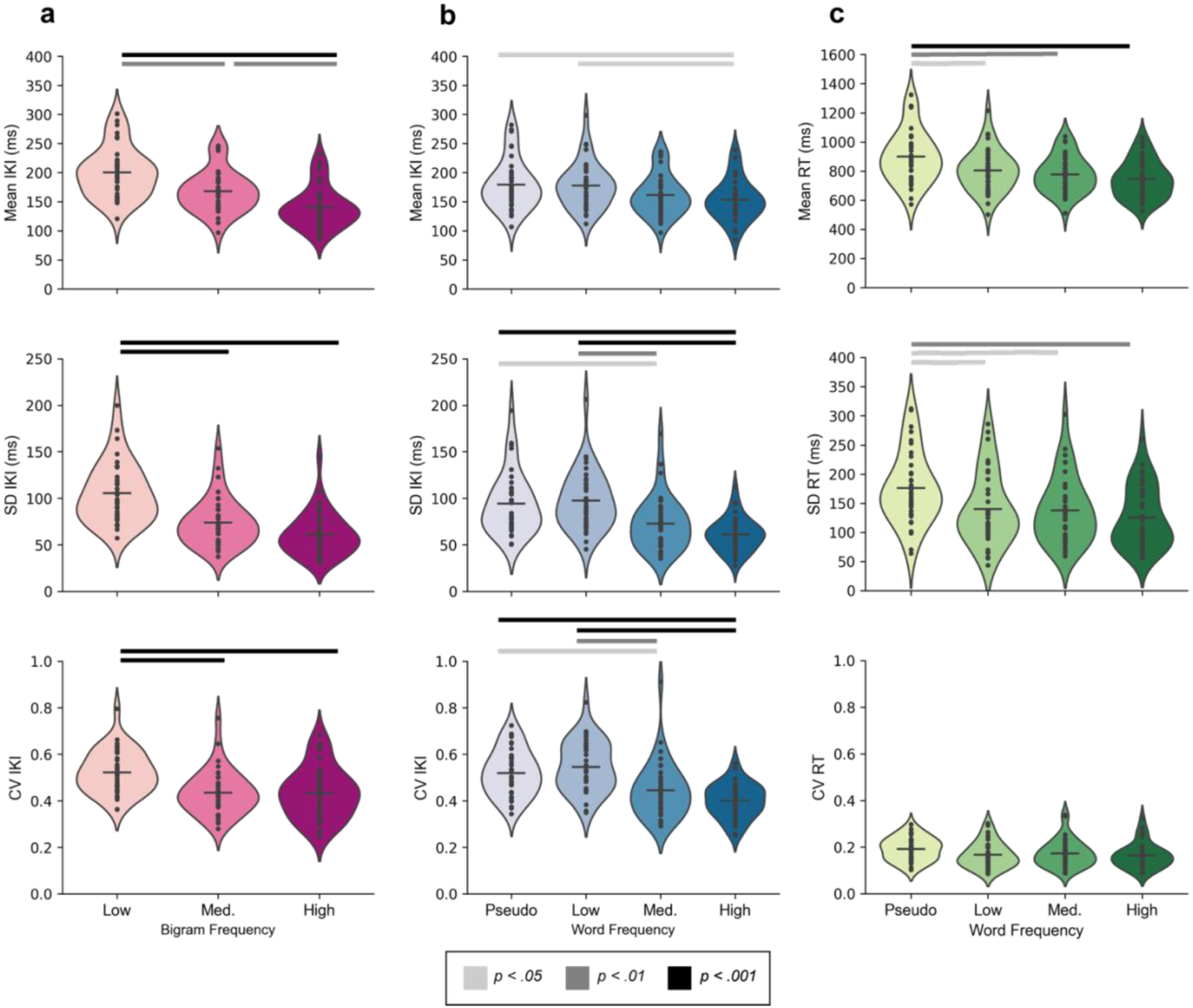
Metrics of interest are shown for bigram frequency IKI, word frequency IKI, and word frequency RT categories (n = 37). Mean, SD, and CV are presented in the top, middle, and bottom rows, respectively. (a) Mean IKI was smaller with increasing bigram frequency across all categories (top). IKI SD and CV were significantly greater for low compared to medium and high frequency bigrams (middle, bottom). (b) Mean IKI differed significantly between pseudo and high frequency words and between low and high frequency words (top). SD and CV IKI for pseudo and low frequency words differed from medium and high frequency words (middle, bottom). (c) Mean RT was significantly longer for pseudo words than all other categories (top). SD RT showed a similar pattern to Mean RT (middle), but CV RT (bottom) showed no significant differences across word frequency categories. The bottom legend indicates p-value thresholds for post-hoc pairwise comparisons.

To evaluate how prior exposure affects trial-to-trial variability in keypress timing, we calculated the within-participant SD IKI for each bigram and word frequency category (Figure 2a, middle row). SD IKI increased with decreasing bigram frequency [(*F*(2,72)=103, *p*=4.7e-18, η^2^_G_=.32] indicating there was greater motor variability when typing less common bigrams. Post-hoc t-tests revealed low frequency bigrams (106±32ms) had significantly higher SD IKIs than medium (74±26ms) and high frequency bigrams (61±23ms) [*p’s*<.001], however the difference between medium and high frequency bigrams was not significant. SD IKI also increased with decreasing word frequency [(*F*(3,108)=36.8, *p*=1.9e-16, η^2^_G_=.22] indicating there is greater motor variability when typing less common strings. SD IKIs for pseudo (97±33ms) and low frequency words (97±31ms) were significantly larger than those of medium (73±29ms) and high frequency words (61±18ms) [*p’s*<.05].

We calculated within-participant CV IKI for each bigram and word frequency category to evaluate trial-to-trial variability proportional to typing speed (Figure 2a, bottom row). Similar to SD IKI, CV IKI decreased with increasing sequence exposure across bigram [(*F*(2,72)=24.1, *p=*9.8e-09, η^2^_G_=.15] and word frequencies [(*F*(3,108)=29.7, *p*=4.4e-14, η^2^_G_=.25]. Post-hoc t-tests showed low frequency bigrams (0.52±0.09ms) had greater CV IKIs than medium (0.44±0.10ms) and high frequency (0.43±0.11ms) bigrams [*p*<.01]. CV IKIs for both pseudo (0.52±0.99ms) and low frequency (0.54±0.11ms) words were greater than medium (0.44±0.12ms) and high frequency words (0.4±0.07ms) [*p’s*<.05]. These data indicate motor variability differences across levels of prior exposure are still present when adjusting proportionally for mean differences in speed.

Mean RT (latency to first key press) increased with decreasing word frequency [(*F*(3,108)=100.9, *p*=3.3e-31, η^2^_G_=.14] (Figure 2b, top). Mean RTs for pseudo words (899±179ms) were significantly longer than medium (777±126ms) and high frequency words (744±129ms) [*p’s*<.01]. SD and CV RT were also calculated to assess differences in reaction time variability across levels of word exposure. SD RT increased with decreasing word frequency [(*F*(3,108)=12.2, *p*=5e-06, η^2^ =.10] (Figure 2b, middle). Similar to mean RT, the pseudo word SD RT (176±61ms) was larger than medium (138±56ms) and high (125±52ms) frequency words [*p’s*<.05]. While there was a main effect of word frequency on CV RT [(*F*(3,108)=3.1, *p*<.031, η^2^_G_=.04], no post-hoc comparisons were significant (Figure 2b, bottom). These patterns show that RT is sensitive to sequence prior practice and that RT variability remains relatively constant when considered proportional to mean RT. Thus, while within sequence variability differed across levels of prior exposure, initiation RTs did not.

### Error rates across levels of prior sequence exposure

As only correct trials were used for the aforementioned comparisons, the numbers of included trials in each category could differ. This was not the case. A comparison of total incorrect trial count across bigram and word categories (Figure 3) showed no significant differences [Bigram: *F*(2,72)=2.59, *p*=.10, η^2^_G_=.02, Word: *F*(3,108)=2.26, *p*=.10, η^2^_G_=.02], indicating participants maintained equivalent accuracy across all frequency categories.

**Figure 3:**
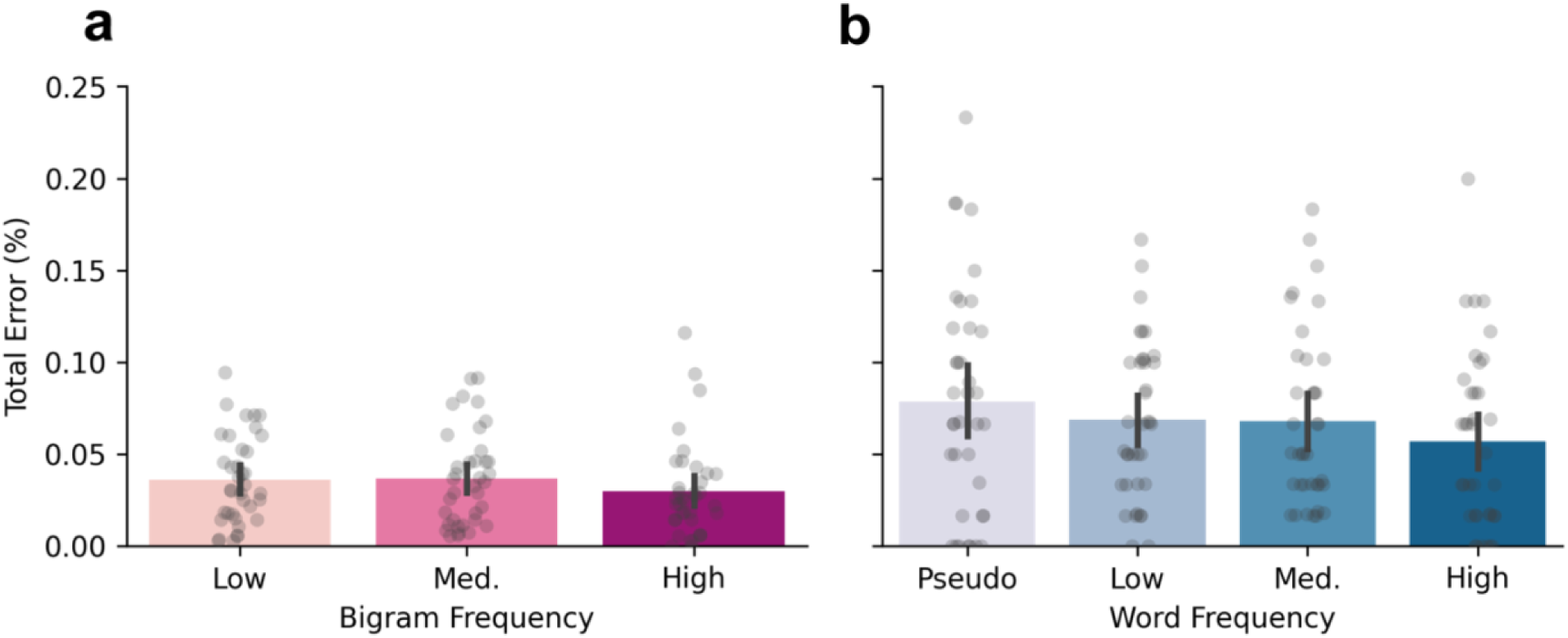
Total error rates (%) for all participants (n = 37) across bigram frequency categories (a) and word frequency categories (b). Grey dots represent individual participant averages. While high frequency bigrams and words tended to have lower error rates, there were no significant differences between categories.

### Individual differences in typing speed and variability

To assess whether our task was sensitive to individual differences, we assessed the correlation of Mean, SD, and CV IKI values between all bigram frequency categories (Figure 4a).

**Figure 4:**
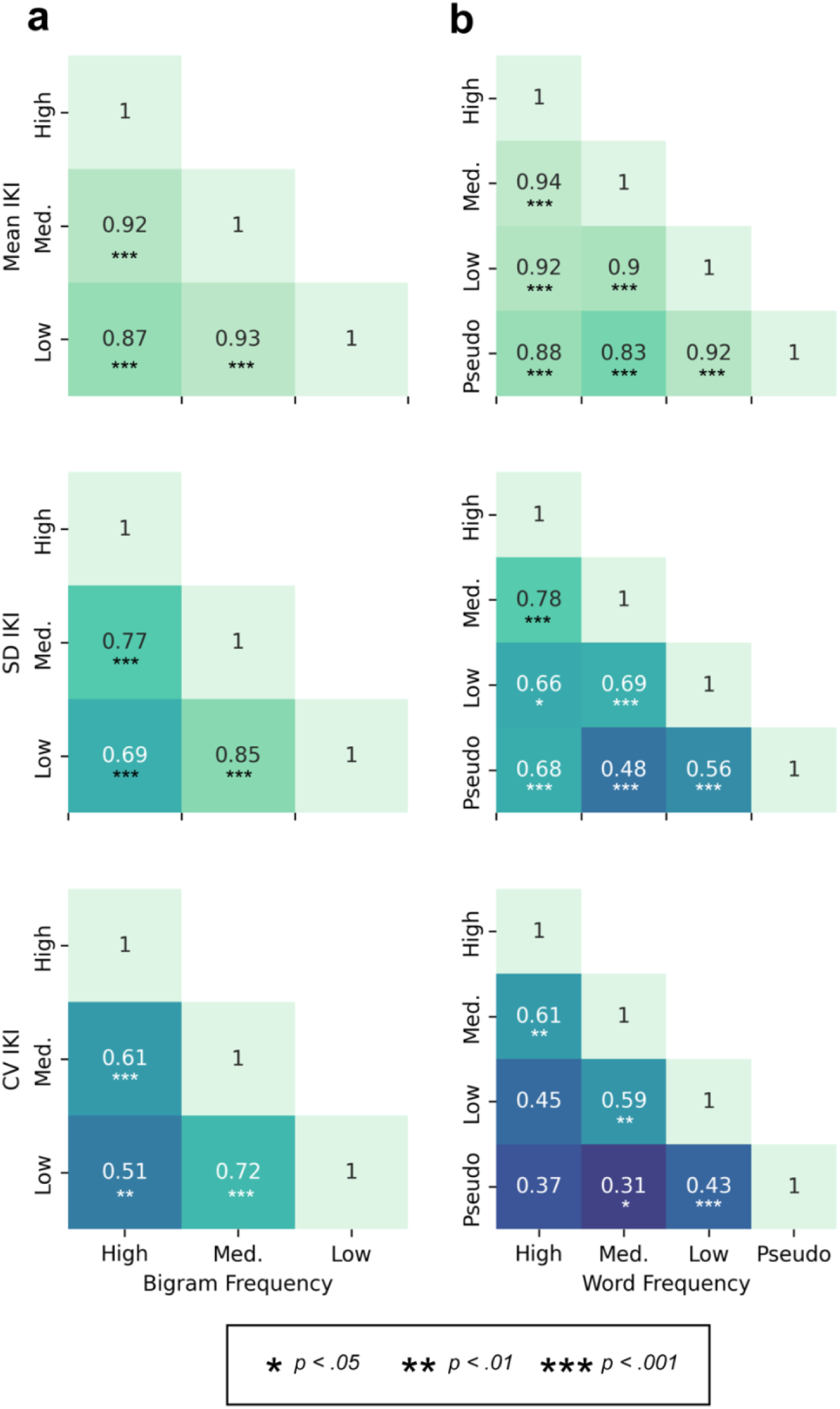
Pearson correlation coefficients (Bonferroni corrected) for comparisons of Mean (top), SD (middle), and CV (bottom) IKI values across bigram (a) and word frequency (b) categories (n = 37). All correlations were significant except for CV IKI pseudo vs high and low vs high word frequencies indicating robust individual differences in performance.

Individuals’ scores were correlated across frequency categories for all metrics. IKI metrics were also compared across all word frequency categories (Figure 4b). Mean and SD IKI yielded positive correlations across all comparisons *(r*’s > .47, *p*’s < .05). CV IKI showed similar results to SD IKI, however pseudo and low vs. high word frequency comparisons were not significant (all other *r*’s > .51, *p*’s < .05). Overall, this suggests our task is sensitive to individual differences in typing speed and variability.

To assess the effect of baseline typing skill on individual differences, mean IES (collected during the pre-task Turbo Typing assessment) was compared with all metrics collected during the 5- character typing task (see Table 1). Mean IES (416±289ms) did not correlate with any of the metrics calculated in our task (Figure 5a; *p’*s > .07). To confirm this finding was not due to the differing characteristics of the two tasks, global mean, SD, and CV IKI were calculated from all alphabetic keypresses during the pre-task Turbo Typing assessment and compared with the same metrics collected from our 5-character typing task (calculated irrespective of frequency category). All metrics correlated between the two tasks [Mean IKI: r(35)=0.83, *p*=1.4e-06; SD IKI: r(35)=0.76, *p*=5.3e-05; CV IKI: r(35)=0.75, *p*=8.9e-09] (Figure 5b). This suggests IKI metrics capture properties of automaticity that are unrelated to conventional typing skill.

**Figure 5:**
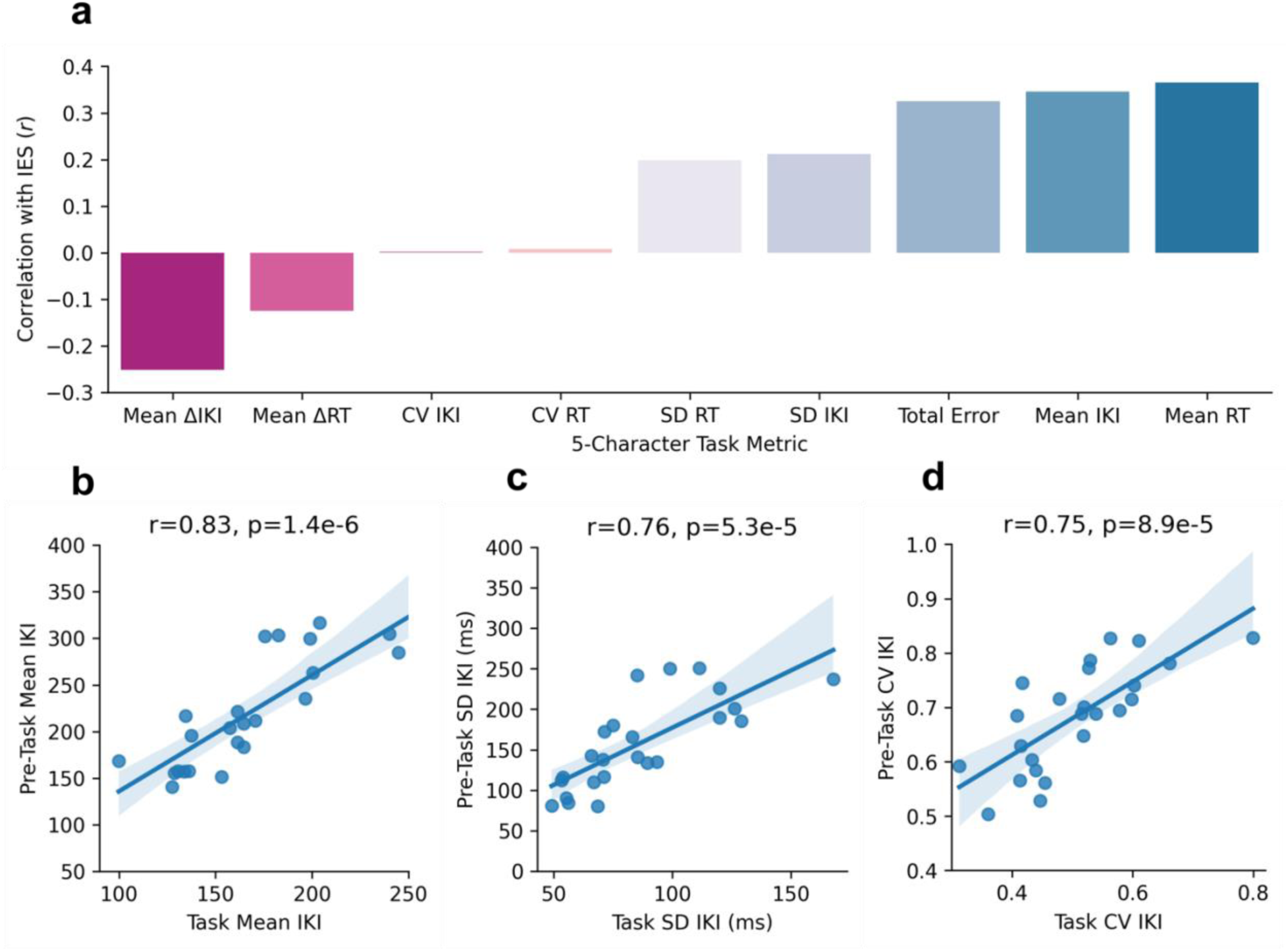
(a) Pearson correlation coefficients from comparisons between IES calculated from the pre-task typing assessment and all collected metrics from the 5-character typing task (n = 24). No metrics correlated significantly with IES. Global (across all frequency categories) mean (b), SD (c), and CV (d) IKI were significantly correlated between the pre-task Turbo Typing assessment and 5- character task.

### Within task “learning” effects on reaction time and interkeypress interval

Mean IKIs and RTs for the last occurrences of all strings and bigrams were significantly faster than the first occurrences, indicating that participants generally reduced their reaction time and speed throughout the task [Mean IKI: t(36)=6.1, *p*=4.7e-07, *d*=.53; Mean RT: t(36)=4.0, *p*=3.4e- 04, *d*=.34] (Figure 6).

**Figure 6:**
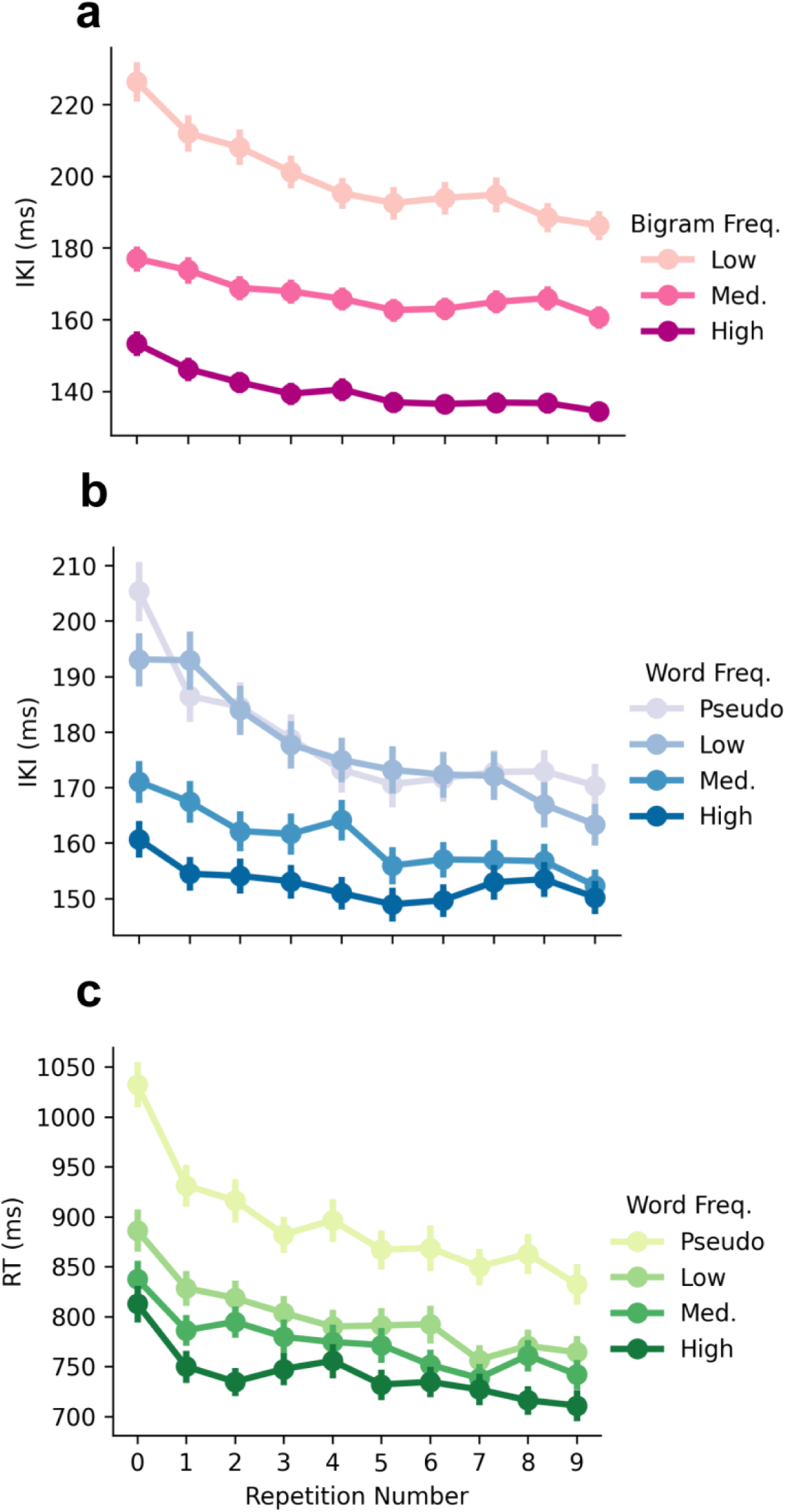
Group mean IKI by bigram frequency (a) and word frequency (b) as well as mean RT by word frequency (c) averaged for each repetition of a given string across the task. Bars represent group standard error. While pseudo and low categories exhibited the largest improvements in speed over the task, all frequency categories showed trends of improvement. Thus, short-term, online improvements can occur even in a highly developed skill.

To determine if prior sequence exposure affects this reduction over the course of the task, change in (Δ) RT and IKI was assessed across consecutive repetitions of the same word. ΔRTs became more negative, reflecting greater improvements in speed, with decreasing word frequency [(*F*(3,108)=13.1, *p*=2.3e-07, η^2^_G_=.16]. RT reductions for pseudo words (-23±16ms) were significantly larger than medium (-9.9±11ms) and high words (-8.9±14ms) [*p’s*<.001] (Figure 6c). ΔIKI became more negative with both decreasing bigram [(*F*(2,72)=24.7, *p*=2.4e-07, η^2^_G_=.19] and word frequency [(*F*(3,108)=12.0, *p*=7.9e-07, η^2^ =.14] (Figure 6a, b). Although the changes across the task were small in magnitude, IKI reductions were significantly larger for low frequency bigrams (-5±4.2ms) than medium (-2±2.2ms) and high frequency bigrams (- 1.9±2.5ms) [*p’s*<.001]. Similarly, reductions were larger for pseudo (-3.9±4.6ms) and low frequency (-4.2±3.7ms) than high frequency (-0.94±2.2ms) words [*p’s*<.01]. These data indicate over the course of repeated trials, RT and IKI became faster for less common sequences than more common ones.

## Discussion

In this study, we introduced a novel approach to studying motor automaticity using typing – a naturalistic and highly practiced behavior. Instead of training novel motor sequences, our task leveraged a daily behavior shaped by lifetime experience. By utilizing linguistic patterns in natural language, we were able to probe how different levels of real-world prior practice influence skilled motor performance. This approach addresses a growing need in the field to examine automaticity within more ethologically relevant behavioral frameworks. While IKI has been used as the main metric to characterize typing in past work (Behmer Jr. & Crump, 2016; Blumenthal-Dramé & McConnell, 2025; Dhakal et al., 2018; Joyce & Gupta, 1990; Logan & Crump, 2009; Pinet et al., 2022; Salthouse, 1984, 1986; Van Waes et al., 2021), our results highlight that IKI variability, rather than speed alone, offers additional insight into motor automaticity. Overall, we found that bigrams are typed faster with increasing prior exposure while variability decreased with prior exposure at both the bigram and string level, even when correcting for differences in speed. However, variability reached a lower limit for the most frequently typed sequences suggesting the presence of a physiological noise floor. While IKI appeared sensitive to levels of automaticity embedded within a sequence, the time to initiate a sequence demonstrated greater sensitivity to the influence of controlled lexical processing.

Specifically, reaction times did not differ across frequency levels for real words, but pseudo words exhibited slower reaction times. Individual differences in typing speed and variability were robust but surprisingly did not correlate with a more conventional measure of typing skill. Finally, we observed trial-to-trial improvements over the course of the task which were greater for less familiar sequences, suggesting online improvements are detectable for automatized components of a skill. Together, these results reveal dissociable contributions of sub-sequence familiarity, whole-sequence structure, and stimulus processing to motor automaticity.

To better understand how prior exposure shapes automated motor performance, we first examined effects at the level of sub-sequences (bigrams). Consistent with previous work (Behmer Jr. & Crump, 2016), we found typing was faster for more frequent bigrams with mean

IKI differing across all bigram frequency categories. To our knowledge, no previous studies of typing have assessed IKI variability in the context of prior sequence exposure. We found variability of typing rhythm, measured via SD and CV IKI, differed between low and medium frequency bigrams, but did not between medium and high frequency bigrams. This suggests speed can continuously decrease in the presence of a limit on variability reduction. There are several possible explanations for the observed variability floor, as motor variability can arise from trial-to-trial differences in the representation of the target location, movement planning, and movement execution (Van Beers et al., 2004).

Well-learned skills, like typing, have been proposed to share mechanistic features with habits through their reliance on stimulus-response (S-R) associations and the caching of action representations paired to corresponding stimuli (Du et al., 2022; Logan, 1979; Yin & Knowlton, 2006). Such caching mechanisms are thought to enable quick and accurate execution by linking stimuli to pre-computed motor outcomes (Haith & Krakauer, 2018). Following this logic, one can infer that automating behavior reduces the time motor computations are exposed to central neural and peripheral noise. Consequently, speed and variability should decrease together as sequences become more familiar. However, our data showed speed and variability diverged for the most familiar words, suggesting more automated actions hit a variability floor, potentially due to physiological and/or peripheral noise during movement execution (Churchland et al., 2006).

Such noise likely comes from multiple physiological sources at the cellular, network, and muscular level (de C. Hamilton et al., 2004; Faisal et al., 2008; Jones et al., 2002; Renart & Machens, 2014).

A related consideration is that the system has tolerance for variability given the task demands. Decreasing variability may not influence typing accuracy beyond a threshold, and the system can continue to speed up as long as accuracy remains unaffected. This suggests speed and variability can be controlled separately and implies automaticity is not controlled through a caching-based system (Cohen & Sternad, 2009). In congruence, growing evidence suggests that habits and automaticity may be dissociable phenomena (Du & Haith, 2023, 2025), and fundamental questions remain concerning what constitutes automatic motor behavior.

We next asked how these effects extend beyond sub-sequences to the level of whole words, where interactions across consecutive keystrokes may introduce additional structure. Our findings suggest typing speed is more sensitive to the naturally occurring frequency of bigrams than words. Unlike the observed differences across all levels of bigram frequencies, only high frequency words were typed significantly faster than low frequency and pseudo words. In terms of variability, words resembled bigram frequencies, exhibiting a floor for medium and high frequency words.

While IKI metrics primarily reflect motor execution, RT provides a complementary measure that captures earlier stages of processing. The observed pattern for mean RT differed markedly from IKI metrics in a manner that suggests RT is sensitive to stimulus processing while IKI is sensitive to motor execution. Mean RTs were slower for pseudo words than the other word frequency categories, which surprisingly did not significantly differ from each other. This dissociation prompted us to consider how stimulus properties, rather than sequence frequency alone, may influence action initiation. Prior studies have found strong correlations between word frequency and RT and posit that less common words require more linguistic processing time (Burgess & Livesay, 1998; Dobbs et al., 1985; Gernsbacher, 1984). However, none of these studies explicitly controlled for bigram frequency leaving open the possibility the initial bigram of a word determines RT. Our task controlled for this possible effect by balancing bigram frequencies across string categories. RT reflects a mixture of component processes including visual processing of the stimulus, initial stages of cognitive processing, motor plan preparation, and execution of the first keypress. The novelty of our pseudo words may have influenced these factors to account for the observed slower action initiation and greater variability. Pseudo words also showed greater SD RT compared to all other frequency categories although this effect was not observed for CV RT.

These findings also have implications for theoretical accounts of automaticity, particularly models based on stimulus–response caching. According to a caching model, we would expect high frequency words, which for many people are typed dozens of times per day, to exhibit faster RTs than low frequency words, which may have never been typed before. However, RT did not differ between these categories. This could be the consequence of RT reaching a floor, but RTs in our task were larger than previous studies (Feldman et al., 2019; Will et al., 2006). It remains possible that RTs for initial bigrams could show differences across word categories and our decision to control for bigram frequency within words effectively eliminated this effect.

To determine whether these timing effects were accompanied by changes in performance accuracy, we next examined error rates across conditions. When comparing error rates across bigram and word frequency, there was a trend of decreasing error with increasing frequency, but surprisingly no significant differences across categories. Only correct trials were used for our time-based metrics, so the observed differences across categories were unlikely due to differences in the number of measurements. Moreover, the pattern of variability did not correspond to error rates.

Beyond group-level effects, we investigated whether these patterns were consistent at the level of individual differences. Notably, IKI correlated positively between all bigram and word frequency categories, and the same was true for variability metrics with a single exception (CV IKI across word frequency). Broadly, this shows individual differences in typing behavior generalize across levels of prior practice, and our collected metrics are sensitive to these individual differences. This aligns with prior research in the cybersecurity field which has shown IKI to be sensitive to individual differences for the potential usage of typed password rhythm as a security biometric (Gaines, R. Stockton et al., 1980; Feldman et al., 2019; Will et al., 2006).

Our data also suggest variability is more sensitive to individual differences at the bigram level than at the word level. This may be a consequence of having a greater number of bigram than word measurements in our study, in this case four times more bigram than word measurements. However, this distinction also raises the possibility that variability at different scales reflects separable underlying mechanisms. Alternatively, variability across bigram categories may be more sensitive to individual differences because it represents lower-level physiological and biomechanical properties with greater intrinsic stability. Conversely, variability at the word level may reflect central motor planning mechanisms and lexical processing that are more adaptable across contexts and less consistent across sequence prior practice (Dhawale et al., 2017).

Assessing these individual differences in future investigations may help identify and distinguish the mechanistic origins and neural circuits supporting variability regulation in linguistic processing and motor performance. Moreover, comparing variability across individuals with differing skill levels can reveal which features of behavior support performance improvements, thereby clarifying how motor automaticity emerges and is maintained in naturalistic tasks.

We next assessed how these individual differences relate to conventional measures of typing skill. Typing skill, as measured by IES in the Turbo Typing assessment, did not correlate with speed or variability metrics derived from our 5-character typing task. This suggests that commonly used measures of skill, which capture the trade-off between speed and accuracy, may be largely unrelated to the low-level motor features commonly attributed to automaticity. Consistent with this interpretation, error rates within our task were not significantly associated with sequence frequency or prior practice, whereas both speed and variability were. These findings suggest that error may be more informative for characterizing practical skill, while speed and variability may better capture the degree of motor automaticity. Notably, overly automated behaviors can also produce errors (Toner et al., 2015), indicating that motor skill and the degree of automaticity may not be tightly coupled. Prior work has shown that typing skill correlates with sensitivity to bigram and trigram frequency, with more skilled typists exhibiting larger speed gains as sequence frequency increases (Behmer Jr. & Crump, 2016). However, in that study skill was defined solely by typing speed. Together, these findings suggest that separately assessing error and speed/variability may provide a more informative framework for studying motor automation (Dhakal et al., 2018).

To confirm this dissociation between conventional typing skill and measures of automaticity was not task-specific, we compared IKI metrics between the two independent typing assessments. When collapsed across frequency categories, mean, SD, and CV IKI from the pre-task assessment were significantly correlated with the corresponding measures from the 5-character task, in contrast to our IES findings. This further confirms the sensitivity of these measurements to individual differences. Overall, these findings suggest that a more granular analysis of IKI speed and variability captures individual differences that extend beyond broad measures of skill, such as IES, and that generalize across distinct task paradigms.

Finally, we examined how performance evolved over repeated trials within the task. Less practiced sequences (both bigram and string) exhibited reductions in IKI and RT over the course of repeated trials. This was not the case for high frequency categories, ruling out a general influence of equipment or task design elements which would be expected to influence all categories. These data indicate that even for a motor skill as well developed as typing, there remains room for improvement when producing less practiced sequences. Within-session reductions of reaction and execution time are often used as markers of early motor learning (Kami et al., 1995; Nissen & Bullemer, 1987; Posner & Keele, 1968; Sakai et al., 2003). Given our task measured performance of a previously learned everyday behavior, it is unclear whether the observed within-session improvements could be considered early-stage motor learning.

Future work can compare typing with a novel sequence learning task to elucidate whether common processes support within-session improvements in performance.

## Conclusion

In summary, we developed a novel behavioral approach to study naturally developed motor automaticity in humans using keyboard typing. By leveraging real-world variation in word and bigram frequencies, we demonstrate that prior exposure systematically influences typing speed and temporal variability, with a variability limit for the most familiar sequences, suggesting a physiological noise floor. Notably, individual differences in these metrics were robust and dissociable from conventional measures of typing skill, underscoring the sensitivity of our approach. We also observed improvements throughout the task for the least familiar sequences, suggesting certain types of performance gains are possible even for well-established and automated skills. Broadly, our findings demonstrate that examination of naturalistic behaviors offers a powerful means of probing late-stage motor skill performance. They also establish a framework for quantifying individual differences in motor automaticity that future studies can directly relate to mechanistic measures, such as neuroimaging or neurophysiological recordings, in both healthy and clinical populations.

## Declarations

### Data availability

Data will be made publicly available upon publication at this repository: https://doi.org/10.5061/dryad.51c59zwrp. This study was not preregistered.

### Code availability

Code will be made publicly available upon publication at this repository: https://doi.org/10.5061/dryad.51c59zwrp.

### Ethics approval

The study was approved by the university’s ethics committee.

### Consent to participate

Consent for publication was obtained from all participants.

### Consent for publication

Consent for publication was obtained from all participants.

### Author contributions

R.R. and I.G. conceptualized the study and designed the experiment. R.R. and M.G. collected the data. R.R. performed data analysis and visualization. I.G., M.B.B., and E.A.W. contributed to methodology and supervision. R.R. and I.G. wrote the original draft of the manuscript. All authors reviewed and edited the manuscript and approved the final version.

## Acknowledgements

We are grateful to the members of the Action Control Lab for insightful discussions, methodological feedback, and support during this project. We especially thank Michelle Marneweck for her insightful feedback on the manuscript alongside Cris Niell and Emily Sylwestrak for assistance with experimental design and study development.

